# Home literacy environment mediates the relationship between socioeconomic status and white matter structure in infants

**DOI:** 10.1101/2021.11.13.468500

**Authors:** Ted K. Turesky, Joseph Sanfilippo, Jennifer Zuk, Banu Ahtam, Borjan Gagoski, Ally Lee, Kathryn Garrisi, Jade Dunstan, Clarisa Carruthers, Jolijn Vanderauwera, Xi Yu, Nadine Gaab

## Abstract

The home literacy environment (HLE) in infancy has been associated with subsequent pre-literacy skill development and HLE at pre-school age has been shown to correlate with white matter organization in tracts that subserve prereading and reading skills. Furthermore, childhood socioeconomic status (SES) has been linked with both HLE and white matter organization. It is also important to understand whether the relationships between environmental factors such as HLE and SES and white matter organization can be detected as early as infancy, as this period is characterized by rapid brain development that may make white matter pathways particularly susceptible to these early experiences. Here, we hypothesized (1) an association between HLE and white matter organization in pre-reading and reading-related tracts in infants, and (2) that this association mediates a link between SES and white matter organization. To test these hypotheses, infants (mean age: 9.2 ± 2.5 months, N = 18) underwent diffusion-weighted imaging MRI during natural sleep. Fractional anisotropy (FA) was estimated from the left superior longitudinal fasciculus (SLF) and left arcuate fasciculus using the automated fiber-tract quantification method. HLE was measured with the Reading subscale of the StimQ and SES was measured with years of maternal education. Self-reported maternal reading ability was also quantified and applied to all statistical models to control for confounding genetic effects. The Reading subscale of the StimQ positively related to FA in left SLF and mediated the association between maternal education and FA in the left SLF. Taken together, these findings underscore the importance of considering HLE from the start of life and may inform novel prevention and intervention strategies targeted at low-SES families to support developing infants during a period of heightened brain plasticity.

## 1 Introduction

Learning to read is a primary goal of elementary education and is associated with subsequent academic outcomes, vocational success, and even health outcomes (Sanfilippo et al., 2020). However, before a child begins formal reading instruction, they develop pre-literacy skills that serve as a foundation for learning to read (de Jong and van der Leij, 1999; Georgiou et al., 2008; Scarborough, 1998; Schatschneider et al., 2004). Some of these developmental milestones are acquired through the child’s home literacy environment (HLE), which generally comprises the interactional extent of shared reading experiences (e.g., duration of parent-child reading) and quality or quantity of reading resources in the home (e.g., number of children’s books). For instance, the frequency and quality of HLE has been shown to aid in children’s pre-literacy skill development, which includes phonological awareness (Frijters et al., 2000), print knowledge (Levy et al., 2006), and oral language skills (Burgess et al., 2002; National Early Literacy Panel, 2008; Storch and Whitehurst, 2002). Predominantly, links between HLE and literacy have been examined in children closer to the time of formal reading instruction (Scarborough and Dobrich, 1994).

Although studies examining HLE in infancy and toddlerhood are rarer, these too have reported that HLE contributes to the prediction of pre-literacy and language outcomes. For instance, infants’ HLE, as measured by parent-informed reports, has been associated with receptive vocabulary during infancy and has been shown to be prospectively associated with expressive vocabulary during toddlerhood (Schmitt et al., 2011). Parent-child shared book reading experience at 8 months also predicted subsequent expressive language abilities at 12 as well as 16 months (Karrass and Braungart-Rieker, 2005). Similarly, receptive and expressive language outcomes at 18 months have been predicted by aspects of shared reading experience (e.g., time spent reading together, children’s interest in reading, maternal questions during shared reading) at 14 months (Laakso et al., 1999) and 10 months (Muhinyi and Rowe, 2019). In children aged 14-30 months, parent-child book reading interactions (specifically quantity of parent utterances) were associated with subsequent receptive vocabulary during 2nd grade and reading comprehension at the end of 3rd grade (Demir-Lira 2015). Also, qualitative aspects of parent-child reading interactions (i.e., caregiver conversational input that elaborates on the text in the book) at 24-months was associated with child receptive vocabulary at roughly 5 years (Malin et al., 2014). The evidence supporting the relationship between HLE during infancy and pre-literacy and language skills later in childhood underscores the importance of considering HLE from the earliest stages of life. However, it is important to note that these observed links between HLE and literacy skill have historically been of modest effect sizes (Bus et al., 1995; Scarborough and Dobrich, 1994) and various studies have suggested that literacy development may be attributed largely to genetic heritability (Hart et al., 2021; van Bergen et al., 2016). Nevertheless, these links between HLE and subsequent reading outcomes are consistent, which suggests that HLE warrants careful investigation of its mechanisms in the context of language and reading development.

In an effort to identify the underlying mechanisms that could explain the link between HLE and subsequent reading outcomes, a few studies have started to examine the brain characteristics related to HLE. Among preschool-age children, HLE was associated not only with better language and pre-literacy skills, but also with white matter organization in pathways thought to be important for reading, including superior longitudinal fasciculus (SLF), arcuate fasciculus (AF), and corpus callosum (Hutton et al., 2020). Similarly, parent-child conversational interactions, which have been subsumed under HLE (Schmitt et al., 2011), were related to fractional anisotropy (FA) in left SLF and left AF in preschool- and kindergarten-age children (Romeo et al., 2018) and to functional connectivity among posterior temporal brain regions in infants (King et al., 2021). Overall, emerging evidence suggests that some of the key characteristics of a high-quality HLE are associated with alterations in left hemisphere white matter tracts at pre-school age. However, the relationship between HLE and white matter structure has not been investigated in children younger than preschool-age. This represents a critical gap as the first two years of life are marked by rapid changes in FA (Geng et al., 2012), potentially making neural circuitry particularly amenable to experiential input during this period (Tau and Peterson, 2010).

It is important to note that the relationship between HLE and brain structure can also be contextualized as part of a larger model relating SES to language outcomes. Such a model has behavioral support in that SES is associated with HLE and this association mediates the relation between SES and pre-literacy outcomes, including phonemic awareness and vocabulary knowledge (Foster et al., 2005), as well as literacy outcomes, including word-level literacy and reading comprehension (Hamilton et al., 2016). However, this certainly does not imply that low HLE and low academic outcomes always accompany low SES (e.g., Christian et al., 1998). Incorporating brain measures, Noble and colleagues postulate that a child’s linguistic environment mediates the influence of SES disparities on language-supporting brain regions, and these brain regions in turn mediate relations between the linguistic environment and subsequent language abilities (Noble et al., 2012). While this model has been tested in older children (5-9 years) and in the context of language (Merz et al., 2020), we propose that an analogous model could apply to literacy and investigated this in infants specifically. In such a model, HLE would mediate the association between SES and brain areas or pathways subserving pre-literacy skills such as the SLF and AF. In older children, the white matter tracts associated with childhood SES include those thought to subserve reading, such as the left SLF (Gullick et al., 2016), left AF (Vanderauwera 2019), left inferior longitudinal fasciculus (Ozernov-Palchik et al., 2018), and left uncinate fasciculus (Vanderauwera et al., 2019). However, to our knowledge, no study (in children of any age) has examined whether HLE mediates the relationship between SES and brain structure. Furthermore, while other brain measures of brain structure and function have been related to SES in infancy (Betancourt et al., 2016; Hanson et al., 2013; Turesky et al., 2019), it remains unclear whether white matter structure relates to SES at this age. This is an important gap to fill to better understand the developmental timeline of brain-SES relations.

The aim of the present study is to examine the relationship between the home literacy environment and white matter organization in infancy. Accordingly, diffusion tensor imaging (DTI) was performed to examine white matter tracts previously associated with HLE in preschool-age children and thought to be important for language and literacy, namely, left SLF and left AF (Hutton et al., 2020). Further, to test the model by Noble and colleagues (Noble et al., 2012), we also investigated whether HLE mediated an association between a prominent aspect of reported SES (i.e., maternal education) and FA in these tracts. Importantly, parental reading ability can contribute to children’s reading ability directly through genes and indirectly through the home literacy environment (Friend et al., 2009, 2008). To control for potential genetic confounds in the present study, we incorporated self-reported maternal reading ability into all models, which can serve as a proxy for genetic transmission (Hart et al., 2021). Ultimately, these inquiries have implications for literacy interventions for developing infants.

## 2 Methods

### 2.1 Participants and Study Design

Infants examined in this study were part of a larger NIH-funded longitudinal investigation of brain, language, and pre-literacy development among children from infancy to school-age (NIH–NICHD R01 HD065762). While the larger study tracks over 150 infants, only 70 of these had diffusion data, and of these, only 38 had HLE and SES datasets of interest. After quality control procedures (see below for details), eighteen infants were included in the final analysis.

Families were recruited from the Greater Boston Area through the Research Participant Registry within the Division of Developmental Medicine at Boston Children’s Hospital, as well as through ads and flyers disseminated in local newspapers, schools, community events, and social media. Families were invited for participation at the time of their child’s infancy with anticipation of continued participation at future developmental time points. Infants completed brain MRI scan sessions and their attending parent(s) completed questionnaires pertaining to the children’s medical history, home environment, and socioeconomic context (please see Table 1 for full demographic details).

**Table 1.**
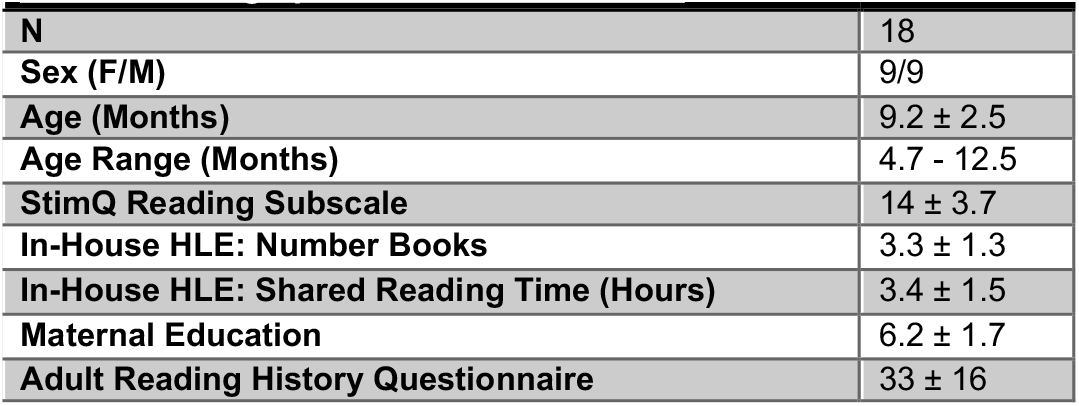
Demographic and Behavioral Data

| **Table 1. Demographic and Behavioral Data** |  |
| --- | --- |
| **N** | 18 |
| **Sex (F/M)** | 9/9 |
| **Age (Months)** | 9.2 ± 2.5 |
| **Age Range (Months)** | 4.7 - 12.5 |
| **StimQ Reading Subscale** | 14 ± 3.7 |
| **In-House HLE: Number Books** | 3.3 ± 1.3 |
| **In-House HLE: Shared Reading Time (Hours)** | 3.4 ± 1.5 |
| **Maternal Education** | 6.2 ± 1.7 |
| **Adult Reading History Questionnaire** | 33 ± 16 |

Participating infants were screened for neurological and sensory impairments, contraindications for MRI evaluation (e.g., metal implants), and premature birth; infants screening positive for any of these criteria were excluded from the study. All children included were from English-speaking families and were born at gestational week 37 or later. Anatomical T1-weighted MRI scans (please see parameters below) were reviewed for clinical abnormalities by a pediatric neuroradiologist at Boston Children’s Hospital, and no participating infants exhibited any malignant brain features. This study was approved by the Institutional Review Board of Boston Children’s Hospital (IRB-P00023182). Informed written consent was provided by each participating infant’s parent(s).

### 2.2 Environmental Characteristics

An inventory of each infant’s home literacy environment (HLE) was gathered from their parents using the StimQ Cognitive Home Environment (infant version), a validated parent report measure that includes four subscales: Availability of Learning Materials, Reading, Parental Involvement in Developmental Advance, and Parental Verbal Responsivity (https://med.nyu.edu/departments-institutes/pediatrics/divisions/developmental-behavioral-pediatrics/research).

The Reading subscale (henceforth termed “StimQ-Reading”), which was the only subscale used in subsequent analyses, instructs caregivers to respond to 15 questions about parent-child shared reading-related activities and the reading materials used. The response scale ranges from 0 to 19 (Table 1).

Socioeconomic status (SES) was measured using maternal education, consistent with previous neuroimaging studies on SES (Betancourt et al., 2016; Brito et al., 2016; Lawson et al., 2013; Merz et al., 2018; Noble et al., 2015; Ozernov-Palchik et al., 2018). Maternal education was coded on an 8-point ordinal scale, with “1” indicating less than 12 years of formal education (less than high school or equivalent), and “8” indicating 20 or more years of formal education (graduate or professional degree; Table 1).

### 2.3 Self-Reported Maternal Reading Ability

Each mother completed the Adult Reading History Questionnaire (ARHQ), which is designed to measure the risk of reading disability in adults by asking questions about the individual’s reading history and current reading habits. Biological mothers answered questions assessing their past and current reading frequency, reading speed, and difficulty reading and spelling, as well as about having to repeat grades or courses, attitudes toward school, and whether they received extra help with learning to read. This questionnaire has demonstrated validity in predicting reading skill (Lefly and Pennington, 2000), making it suitable to use in this study as a measure to control for genetic transmission of reading abilities from mother to child (please see Introduction). All responses were given on a Likert scale and summed for each mother and lower values indicate greater reading ability (Table 1).

### 2.4 Neuroimaging Data Acquisition

Neuroimaging data were acquired while all the infants were naturally sleeping, without sedation, using an established infant neuroimaging protocol (Raschle et al., 2012; T. K. Turesky et al., 2021). Diffusion-weighted and structural T1-weighted images were acquired on a 3.0 Tesla Siemens MRI scanner with a standard Siemens 32-channel radio frequency head coil. One parent remained in the MRI room with the infant for the duration of the scan, in addition to a researcher who stood by the bore to monitor changes in the infant’s sleeping state and potential motion. Structural T1-weighted whole-brain multi-echo magnetization-prepared rapid gradient-echo sequences with prospective motion correction (mocoMEMPRAGE) were acquired for each participant with the following parameters: repetition time (TR) = 2270 ms; echo time (TE) = 1450 ms; acquisition time (TA) = 4.51 min; flip angle = 7°; field of view = 220 x 220 mm; voxel size = 1 x 1 x 1 mm3; 176 slices; in-plane GRAPPA acceleration factor of 2. Diffusion echo planar images were acquired using the following parameters: TR = 4600 ms; TE = 89 ms; TA = 5.36 min; flip angle = 90°; field of view = 256 x 256 mm; voxel size: 2 x 2 x 2 mm3; 64 slices; in-plane GRAPPA acceleration factor = 2; slice-acceleration (SMS/MB) factor = 2; partial Fourier encoding = 6/8, 30 gradient directions of b = 1000 s/mm2, 10 non-diffusion-weighted volumes of b = 0 s/mm2 acquisitions, 1 phase-encoding anterior-to-posterior (AP) volume, 1 phase-encoding posterior-to-anterior (PA) volume.

### 2.5 Diffusion-Weighted Imaging Processing

Raw DWI data in DICOM format were converted to NRRD file format using the DWIConvert module of Slicer4 (https://www.slicer.org/) and submitted to DTIPPrep to identify volumes with excessive motion artifacts (2 mm translation threshold and 0.5° rotation threshold). Artifactual volumes were removed prior to subsequent processing while non-artifactual DWI data were corrected for susceptibility distortions using FSL’s topup module. Eddy currents were corrected and all gradient volumes were realigned to the b0 volumes using FSL’s eddy module (https://fsl.fmrib.ox.ac.uk/fsl; Smith et al., 2004). Tensor fitting was then performed using mrDiffusion in the VISTALab diffusion MRI software suite (https://vistalab.stanford.edu/). FA quantification was conducted along the trajectory of each tract based on eigenvalues from the diffusion tensor estimation (Basser et al., 1994).

### 2.6 Automated Fiber Quantification

The Automated Fiber Quantification (AFQ) software package was used to quantify white matter organization in left superior longitudinal fasciculus (SLF) and left arcuate fasciculus (AF; https://github.com/yeatmanlab/AFQ; Yeatman et al., 2012). The SLF is a tripartite tract and mainly connects parietal to frontal areas (Yagmurlu et al., 2016). With the methods we employed, all segments were captured as a single tract (Yeatman et al., 2012). Relevant to the inquiries posed in this study, one segment (SLF III) constitutes the anterior segment of the dorsal language pathway (Catani and Dawson, 2017), which has a role in fluency and naming (Ivanova et al., 2021). It connects supramarginal gyrus to inferior frontal gyrus (Yagmurlu et al., 2016), two regions involved in reading and reading-related processes (Eden et al., 2016). The AF constitutes the long segment of the dorsal language pathway (Catani and Dawson, 2017). It connects superior temporal gyrus and inferior frontal gyrus, and is important for naming abilities (Ivanova et al., 2021), word repetition (Sierpowska et al., 2017) and reading (Gullick and Booth, 2015; Thiebaut De Schotten et al., 2014).

Similar to previous studies (Langer et al., 2017; Zuk et al., 2021), whole-brain tractography was computed using a deterministic streamline tracking algorithm, whereby fiber tracking was terminated in instances where estimated FA was below a threshold value of 0.15 and the angle between the last path segment and the next direction was greater than 40°. Region of interest (ROI)-based fiber tract segmentation and fiber-tract cleaning were then employed using a statistical outlier rejection algorithm. FA of each fiber tract was sampled to 100 equidistant nodes. Left SLF and left AF tracts were subsequently inspected visually by two separate raters to verify successful reconstruction (please see Figure 1 for a participant with successful left SLF and left AF reconstructions). Out of an initial 38 infants with StimQ, maternal education, and diffusion data, only 18 infants had successful left SLF reconstructions and only 19 infants had successful left AF reconstructions.

**Figure 1.**
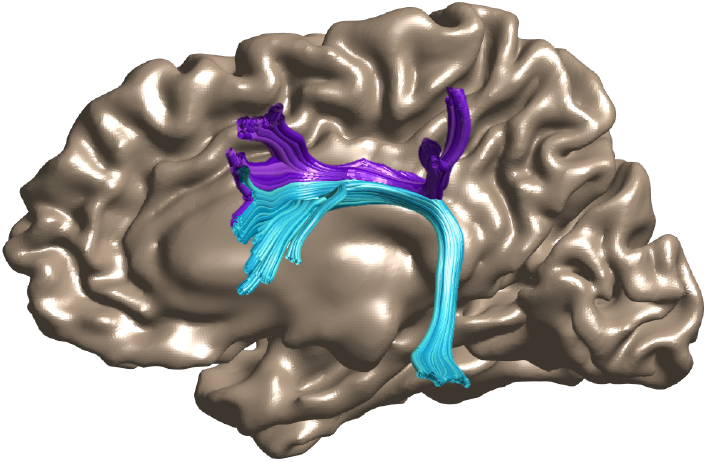
Tract reconstructions for one infant. Left superior longitudinal fasciculus (purple) and left arcuate fasciculus (blue) in one infant are shown for one infant, superimpose on a standard mesh brain, rescaled for a typical infant.

### 2.7 Statistical Analyses

Correlation analyses were conducted among HLE, FA, and SES variables, and where HLE variables correlated with both FA and SES, subsequent mediation analyses were performed. All correlation analyses were conducted in MATLAB. Because the distribution of StimQ-Reading was not normally distributed according to D’Agostino Pearson omnibus normality tests (K2 = 7.18, p <0.05), non-parametric statistics (i.e., Spearman) were used. First, to examine whether HLE and maternal education were related, semipartial correlations between StimQ-Reading (adjusted for infant age at time of MRI scan and self-reported maternal reading ability) and maternal education were computed.

Second, relations to FA for nodes along each tract of interest (i.e., SLF and AF) were done according to partial correlations (adjusting for infant age at the time of scan and self-reported maternal reading ability) for StimQ-Reading scores and semipartial correlations for maternal education. Significance testing was done using non-parametric bootstrapping to simulate the sampling distribution in the general population to ensure proper evaluation of effect sizes (Zuk et al., 2021). This approach involved simulating the population distribution based on partial (for StimQ-Reading) or semipartial (for maternal education) using 5000 replicated samples with replacement from the original sample (Nichols and Holmes, 2002). To correct for multiple tests of correlation within each tract (100 tests), a cluster-based non-parametric permutation method was applied using an adapted version of the AFQ_MultiCompCorrection function in the AFQ suite adjusted to include covariates and Spearman testing. This function estimated the number of contiguous nodes significant at an uncorrected p <0.05 that were needed for a family-wise error correction (FWE) p < 0.05, which differed according to tract and behavior estimate examined. FA estimates of contiguous nodes passing correction for multiple comparison were then averaged for visualization purposes.

Finally, a 10,000-repetition bootstrapped, non-parametric mediation analysis was run using R statistical analysis software, with maternal education as the independent variable, FA as the dependent variable, StimQ-Reading as the mediator, and infant age at time of MRI and self-reported maternal reading ability as confounding variables. The FA values used in the mediation were taken from the contiguous nodes that were significant (after FWE correction) in correlations with StimQ and maternal education. Among these nodes, those with an additional node-wise threshold of p < 0.01 were averaged and submitted to mediation analyses. Effects were deemed significant when 95% confidence intervals did not include 0.

## 3 Results

Among the 18 infants with StimQ data, the overall score on the Reading subscale of the StimQ (henceforth “StimQ-Reading”) was correlated with maternal education (r = 0.48; p <0.05). StimQ-Reading also correlated with FA in the mid-portion of the left SLF between nodes 22 and 34 (out of 100), inclusive, controlling for age at time of scan and self-reported maternal reading ability (r_average_ = 0.61, p_FWE_ <0.05, Figure 2). No other segment of the left SLF or any segment of the left AF showed a significant correlation with the StimQ-Reading subscale after FWE correction.

**Figure 2.**
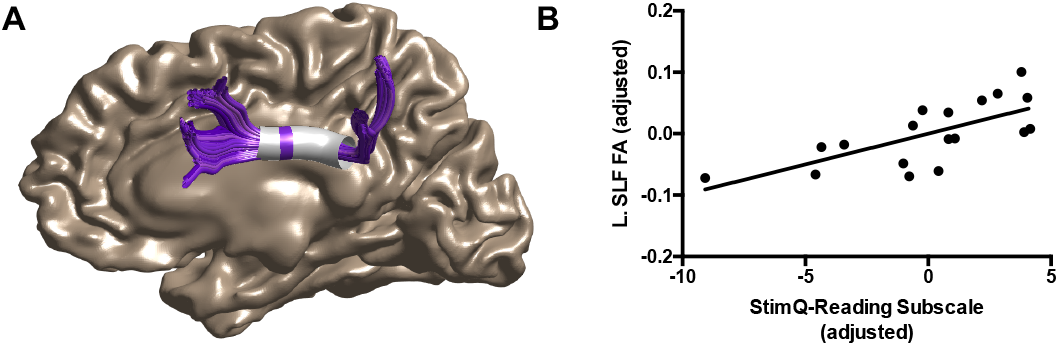
Association between StimQ-Reading Score and FA in left SLF. (A) Nodes exhibiting significant (after FWE correction for multiple comparisons) associations with the StimQ-Reading Score (purple) superimposed on a midsagital slice of one infant. (B) Scatterplot depicting average FA from nodes represented in A in relation to StimQ-Reading Score. Both variables were adjusted for infant age and self-reported maternal reading ability. No outliers were identified according to the ROUT (Q = 1%) or Grubbs ( = 0.05), but results remained significant (with FWE correction) after removal of the participant with the lowest (adjusted) StimQ score.

Maternal education within this group was also strongly correlated with FA of the left SLF between nodes 22 and 33, inclusive, controlling for age of the infant and self-reported maternal reading ability (r_average_ = 0.58, p_FWE_ <0.05, Figure 3). There was also a significant correlation between maternal education and FA of the left SLF at nodes 61-71 (r_average_ = 0.56, p_FWE_ <0.05). No segment of left AF showed any significant correlation with maternal education after FWE correction.

**Figure 3.**
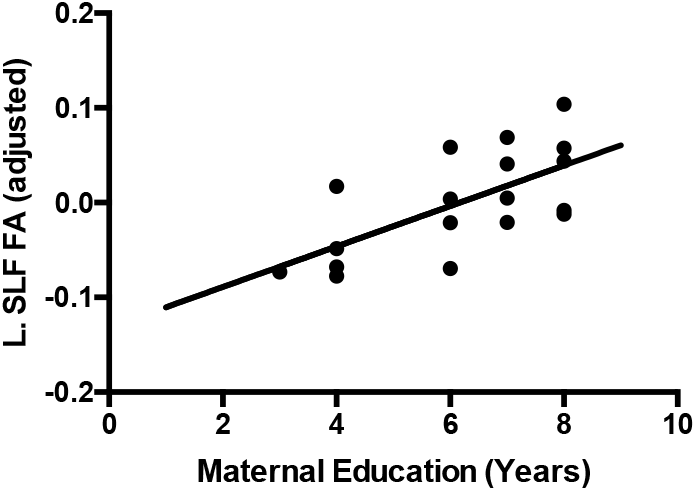
Association between maternal education and FA in left SLF. Maternal education was significantly (after FWE correction for multiple comparisons) associated with FA in left SLF in 12 nodes that overlapped with the nodes associated with StimQ-Reading Score. FA from these nodes were averaged, adjusted for infant age and self-reported maternal reading ability, and depicted on the y-axis of this scatterplot.

Given that StimQ-Reading correlated with maternal education and FA in SLF, a mediation model was constructed to test for an indirect effect of StimQ-Reading on the relationship between maternal education (independent variable) and FA of the left SLF in the cluster of nodes that overlapped in the FA-stimQ and FA-maternal education associations above (dependent variable). Controlling for infant age and self-reported maternal reading ability, StimQ-Reading partially mediated the relationship between maternal education and average FA of the left SLF at nodes 29-32 (direct effect = 0.015, [95% CI = 0.0024-0.030], p <0.05; indirect effect = 0.0095 [95% CI = 0.0014-0.020], p <0.05; indirect/total effect = 0.39; Figure 4), such that the StimQ-Reading score accounted for approximately 39% of the total relationship between maternal education and the left SLF cluster FA.

**Figure 4.**
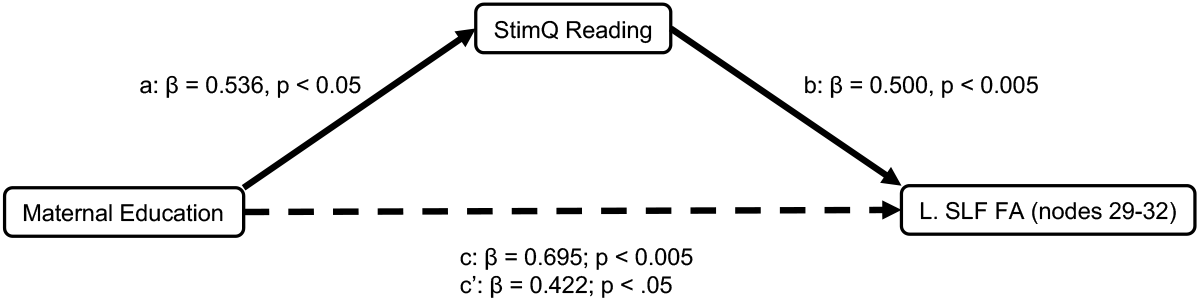
Schematic of the mediation model. Significant standardized regression coefficients are shown between (a) maternal education and StimQ-Reading, (b) StimQ-Reading and the average FA of nodes 22-33 of the left SLF controlling for maternal education, (c) maternal education and left SLF FA, and (c’) maternal education and left SLF FA with StimQ-Reading. The model also included infant age at the time of scan and self-reported maternal reading ability as covariates.

## 4 Discussion

In the present study, we show that the home literacy environment (HLE), as measured with the Reading subscale of the StimQ, correlates with white matter organization in the left superior longitudinal fasciculus and partially mediates the relationship between socioeconomic status and left SLF white matter fractional anisotropy (FA). Along with previous work showing that white matter in infancy relates to later language outcomes (Zuk et al., 2021), these findings offer a neurobiological mechanism to the corpus of behavioral literature linking HLE during infancy and toddlerhood to pre-literacy and language development (Karrass and Braungart-Rieker, 2005; Laakso et al., 1999; Malin et al., 2014; Muhinyi and Rowe, 2019; Schmitt et al., 2011). Further, the finding of an environmental factor relating literacy to white matter organization (independent of familial transmission as measured by self-reported maternal reading ability) complements previous reports showing that familial risk of reading difficulty (which did not relate to HLE in their cohort) was related to white matter organization in infancy (Langer et al., 2017). Taken together, our findings indicate that HLE is associated with white matter organization early in development, prior to prolonged exposure to literacy input. Our main findings were localized to a segment in the mid-portion of the left SLF, which is a tripartite tract that mainly connects parietal to frontal areas (Yagmurlu et al., 2016) and constitutes the anterior segment of the dorsal language pathway (Catani and Dawson, 2017). Further illustrating the importance of the left SLF in literacy, FA in this tract has been shown to correlate with reading outcomes in typically developing children (Hoeft et al., 2011; please note, this study defined left SLF as including left arcuate fasciculus), while children with a family history of dyslexia have greater FA in the left SLF compared with children without a family history of dyslexia at beginning and fluent reading stages (Wang et al., 2017). The combination of our results linking HLE to left SLF anatomy and past findings relating left SLF to literacy outcomes supports behavioral literature linking HLE in infancy to language outcomes (Duff et al., 2015; Laakso et al., 1999; Amber Muhinyi and Rowe, 2019; Schmitt et al., 2011). However, follow-up work examining reading outcomes in the children in this study and causal mediation testing will be needed to confirm the hypothesis that HLE impacts subsequent reading ability via left SLF structure. Broadly, findings in infants of an association between HLE and white matter structure are consistent with work previously done in pre-school age children. Namely, HLE as measured with StimQ-Reading has been associated with white matter organization in the SLF and other literacy-supporting white matter tracts as measured with whole-brain, tract-based spatial statistics (TBSS) (Hutton et al., 2020), as well as with activation in left hemisphere brain regions in or near SLF termini, such as inferior frontal gyrus (Hutton et al., 2017; Powers et al., 2016) and the parietal-temporal-occipital region (Hutton et al., 2015) on story listening and phonological processing tasks. However, in pre-school age children, the association between HLE and left SLF was mainly measured with axial diffusivity (and not FA). Also, HLE was associated with white matter structure in other tracts supporting language and literacy (e.g., arcuate fasciculus and inferior longitudinal fasciculus), in some cases bilaterally (Hutton et al., 2020). In the present study in infants, links were only observed for the left SLF as estimated using a hypothesis-driven, two-tract-specific approach (with the left arcuate fasciculus being the sole other tract examined). Such differences between the two developmental stages, though not directly tested here, would not be surprising, as white matter in all tracts undergoes substantial change between infancy and preschool (Lebel and Deoni, 2018; Reynolds et al., 2019), and it is conceivable that variation in the developmental trajectories of these tracts would make them more or less amenable to environmental input at different developmental stages (Nelson and Gabard-Durnam, 2020). Future longitudinal studies will be needed to determine whether the associations between HLE and white matter structure indeed become more widespread with age. The present study also contributes to the literature on HLE by partially controlling for genetic confounds using self-reported maternal (but not paternal) reading ability. This is important because genetic makeup shared by parent and child contributes to the child’s reading ability directly via genetic inheritance and indirectly via parental reading ability and the home literacy environment (Hart et al., 2021; van Bergen et al., 2016). Past studies using parental characteristics as a proxy for the direct/genetic pathway have largely reported diminished associations between HLE variables and children’s reading when controlling for parental reading ability (van Bergen et al., 2016) or reading-related skills (Puglisi et al., 2017). Interestingly, the one HLE variable to remain significantly associated with reading outcomes after controlling for parental reading ability was number of books in the home (van Bergen et al., 2016), which is the one individual HLE component we identified as related to white matter organization in the left SLF. One study found associations between HLE and child vocabulary after controlling for parental IQ/vocabulary, which the authors describe as a genetic variable; however, this study did not examine the relation between HLE and child outcomes prior to controlling for parental IQ, making it difficult to determine the effect of the genetic confounding factor (Storch and Whitehurst, 2001). As suggested by Hart and colleagues (2021), we have used self-reported maternal reading ability as a proxy for genetic transmission, and have found relations between HLE and FA despite its inclusion, suggesting that HLE relates to white matter organization independent of genetic influences. Our results also speak to the association between SES and brain structure. Though consistently reported (Hanson et al., 2011; Jednorog et al., 2012; Luby et al., 2013; McDermott et al., 2019; Noble et al., 2015, 2012), including for white matter organization (Gullick et al., 2016; Ozernov-Palchik et al., 2018; Vanderauwera et al., 2019), the mechanism through which SES would influence brain structure is rarely examined empirically (for a discussion, please see (Farah, 2017)). One exception is a report by Merz and colleagues (2020), which reported that linguistic input (measured as the principal component of adult words and conversational turns) mediated the relation between parental education and left perisylvian surface area (Merz et al., 2020). Our results complement these findings by identifying indirect effects between maternal education and white matter organization in the left SLF via literacy input and further extend this pathway to children as young as infants. These results also could have important implications for interventions because compared with SES, HLE is a modifiable environmental factor for which interventions can be designed to optimize language and literacy development (Powers et al., 2016). “Reach Out and Read” is one such intervention, which distributes literacy materials to low-SES families in primary care medical clinics, and has demonstrated efficacy in improving language and preliteracy skills in children (Zuckerman, 2009). Concordantly, pediatricians and other clinicians may be aptly situated to educate families on HLE while the child is in their first year of life. Importantly, children with a familiar risk for developing literacy or language difficulties may be especially vulnerable to the effects of rearing in low-SES backgrounds and/or with lower quality HLE, since they would be incurring both genetic and environmental adversity (for discussions of gene-environment interactions related to reading and reading difficulty, please see Hart et al. (2021) and van Bergen et al., (2014)).

This study had five main limitations. The first limitation has to do with our approach to controlling for genetic confounding, which was limited to selfreported maternal reading ability, rather than the reading abilities of both parents. Because children share 50% of genes with one parent, our approach did not control for genetic confounding. Moreover, the Adult Reading History Questionnaire used here is not a direct measure of reading ability, but a reliable correlate (Lefly and Pennington, 2000), and as with direct measurements of reading ability, it may serve as a proxy for genetic transmission (Hart et al., 2021; van Bergen et al., 2016). Second, we did not examine individual components of HLE. While previous work has shown weaker associations between individual HLE components and language outcomes, compared with robust composite associations, additional other work has suggested that individual HLE components may relate differently to outcomes (Burgess et al., 2002; Muhinyi and Rowe, 2019; Storch and Whitehurst, 2001). Third, the infants in this study are from a relatively narrow range of high SES families and it is not clear whether the relations we observed between HLE and white matter organization in the left SLF would persist in infants from low SES families. Some, for example, have suggested that brain-behavior relations may depend on the degree of socioeconomic insufficiency (Brito and Noble, 2014; Turesky et al., 2021). While previous findings of HLE-outcome relations in children from low-income families (Payne et al., 1994) suggest that HLE-brain relations would indeed persist, empirical testing would be needed to confirm this. Fourth, our sample was limited to traditional nuclear families in which infants are raised by two biological parents of opposite sex. It is not entirely clear whether our findings would generalize to nontraditional families (e.g., non-biological parents). Fifth, the sample size represented here was modest. Overall, it might behoove future studies examining relations between HLE and early white matter organization to examine larger sample sizes, a wider SES range, individual HLE components, and genetic data from both parents to more accurately capture genetic confounding variables and to disentangle gene and environmental influences (Hart et al., 2021).

In conclusion, this study examines relationships among HLE, SES, and white matter organization in infancy. Results show that HLE is associated with FA in the left SLF and mediates the link between maternal education and left SLF FA. With these links to white matter organization, this study fills an important gap in behavioral literature linking HLE and SES to neurocognitive outcomes and further evinces the importance of HLE in child development.

## 5 Acknowledgements

This work was funded by NIH–NICHD R01 HD065762, the William Hearst Fund (Harvard University), and the Harvard Catalyst/NIH (5UL1RR025758) to N.G., the Harvard Brain Initiative Transitions Program to T.K.T, the Ruth Taylor Research Fund (Queen’s University) to J.S., and the Sackler Scholar Program in Psychobiology to J.Z. We would like to thank all participating families for their long-term dedication to this study. We are grateful for all additional members of the research team who contributed to data collection and management and quality control, especially Bryce Becker, Danielle Silva, Michael Figuccio, Doroteja Rubez, and Elizabeth Escalante. We also thank Carolyn King for her feedback on the manuscript.

## Notes

### Competing Interest Statement

The authors have declared no competing interest.

